# Effects of environmental setting and diet on the gut microbial ecology of eastern hellbenders (*Cryptobranchus alleganiensis alleganiensis*)

**DOI:** 10.1101/2025.11.24.690264

**Authors:** Chloe Cummins, William Sutton, Taina McLeod, Jason W. Dallas, Mitra Ghotbi, Lluvia Vargas-Gastélum, N. Reed Alexander, Alexander J. Rurik, Dale McGinnity, Sherri Doro Reinsch, Pia Sandonato, Jessica Arbour, Michael Freake, Anthony Ashley, William Ternes, Elizabeth Culp, Joseph Spatafora, Kerry McPhail, Jason E. Stajich, Rebecca Hardman, Donald M. Walker

**Author notes:** Corresponding Author: Donald M. Walker.

## Abstract

**Background:** Eastern hellbenders (*Cryptobranchus alleganiensis alleganiensis*) have undergone substantial population declines throughout their range, leading them to become the focus of increased conservation efforts, including care in zoo and university settings. However, effective implementation of such conservation strategies often relies on a comprehensive understanding of host health, which can be directly influenced by the gut microbiome, yet characterization of gut microbiota often remains overlooked in ex situ conservation facilities. Additionally, effects on the gut microbiome associated with releasing zoo-reared animals into the wild are poorly understood. Therefore, these circumstances make hellbenders an ideal species to examine the relationship between zoo management strategies and gut microbial dynamics.

**Methods:** 16S rRNA sequencing was used to investigate dissimilarities between the gut microbiome of hellbenders in zoo and wild settings and to evaluate the impact of implementing a wild diet in zoo-reared hellbenders. Additionally, the bacterial composition of zoo-released individuals and wild resident hellbenders was compared to examine the response of the gut microbiome upon release into natural habitat. Selected samples were also chosen for ITS1 rDNA sequencing as a preliminary investigation of the hellbender gut mycobiome.

**Results:** Human rearing strongly affected the gut microbiome, leading to reduced bacterial richness as well as differing community structure than wild hellbenders. However, implementation of a wild diet in a zoo setting modulated the microbiome and appeared to be mainly driven by bacterial turnover. Additionally, both bacterial and fungal gut assemblages demonstrated the capacity for restructuring upon release into native habitat to become more reflective of a wild-type microbiome.

**Conclusions:** We completed the first study elucidating the gut microbial composition patterns of hellbenders, across both zoo and wild settings. These results provide an understanding of the potential impacts of conservation populations in zoos on gut microbial communities and also inform headstart programs of the transition of the gut microbiome post-reintroduction to the wild.

## BACKGROUND

Amphibians are the most threatened group of vertebrates [1, 2], with recent estimates indicating that 41% are threatened with extinction [3]. Current decreases in amphibian abundance and diversity have been attributed to a myriad of potential causes, including climate change, emerging infectious diseases, and habitat loss [1, 4, 5]. Further, these declines have stimulated an increase in management strategies, such as headstart programs, for species conservation [6, 7]. However, conservation initiatives require a comprehensive understanding of host health in conservation populations while under human care as well as prior to and following reintroduction into natural habitats for successful translocations. Notably, the gut microbiome is directly linked to and strongly influences host health by aiding in immune system function and conferring defense against pathogens [8–10], but it is frequently overlooked in conservation management programs [8, 11–13].

The assembly of the gut microbiome can be influenced by a variety of intrinsic (i.e., host-mediated) and extrinsic factors, such as diet and the environment [8, 14–19]. The host microbial assemblage is also susceptible to environmental disturbance, and disturbance-induced dysbiosis of the gut microbiome can negatively impact host survival [8–10]. According to the Anna Karenina principle [20], the community composition of dysbiotic microbiomes feature high dispersion, or greater taxonomic variation, in response to disturbances, whereas “healthy” microbiomes are defined by low variation and greater stability. Dysbiosis of the gut microbiome is associated with increased susceptibility to pathogenic infections, chronic morbidity, and decreased host fitness [8, 21–23]. Thus, the need for characterizing the gut microbiome and documenting consequences of deviations from natural microbial communities in species of conservation concern has been colloquially described as a “microbial renaissance” [13].

In wildlife gut microbiome research, the effect of human care on gut microbial communities is a recurring theme; however, research focused on amphibians remains minimal. Though investigating gut assemblages is a critical area of research, it is important to also consider that human rearing may not ultimately reflect the conditions that these animals would be exposed to in their natural environments [12]. Factors associated with ex-situ populations under human care, such as increased stress levels, antibiotic use, lack of exposure to natural habitat, reduced contact with conspecifics, and altered diets, have been shown to influence microbiome composition and function [16, 17, 24–26], resulting in a divergence from wild conspecifics [16, 23, 27, 28]. In many cases, the microbial communities of zoo animals feature reduced taxonomic diversity [23–26, 29] and are often dominated by or enriched with pathogenic taxa that may be indicators of potential disease or poor health [16, 23, 28, 30]. Therefore, to reveal key insights into host health, elucidating the composition and structure of the gut microbial community in zoo-reared organisms is an important consideration for the management of species in headstart programs and other conservation efforts. Comparing the gut microbiomes of individuals in ex-situ programs and wild individuals can guide specific practices to promote colonization by diverse microbiota typically observed in wild counterparts [12, 13, 31].

The concept of “rewilding” has gained interest as a method to promote the formation of healthy or more natural microbiomes in conservation approaches. In the context of microbial ecology, rewilding involves exposing hosts to natural environmental conditions with the goal of promoting the colonization of native resident microbiota [32–34]. In zoos, lab settings, and rescue centers, rewilding is implemented through various techniques to enhance host health, such as providing individuals with more natural diets and exposing them to environmental substrates that can act as microbial reservoirs [31, 33–36]. For example, white-footed mice (*Peromyscus leucopus*) under human care that were fed a wild diet developed gut microbiomes more similar to wild and reintroduced mice than to zoo-reared mice on a standardized diet [36]. Additionally, restructuring of the microbiome has been investigated in translocation and reintroduction efforts to determine whether gut microbiomes associated with ex situ conditions, which may result in depauperate microbial communities, persist or revert to a more natural state upon release to the wild [32, 35, 37]. Incorporating the rewilding of wildlife microbiomes may act as a promising approach to improve repatriation initiatives, particularly for host groups of conservation concern like amphibians.

Eastern hellbenders (*Cryptobranchus alleganiensis alleganiensis*) have experienced considerable population declines across their historic range in recent decades [38–40]. Approximately 75% of historical populations are extirpated, functionally extirpated, or in decline, and extirpations are projected to increase by 20-80% in the coming decades [41]. The proposed primary causes of these declines involve habitat loss and degradation, particularly declining water quality due to sedimentation and river impoundment [41, 42]. Researchers have also observed a demographic shift in age class structure towards older individuals, a pattern indicative of reduced recruitment and potentially lower reproductive success [38, 43, 44]. Similar observations have been made for some eastern hellbender populations in Middle Tennessee [45], and statewide, hellbender populations have experienced an estimated 61% decline since 2000 [40]. In response, many zoos and conservation organizations have established hellbender conservation initiatives, such as propagation, headstarting, and translocation efforts. These programs offer a unique opportunity to investigate the composition and community structure of the gut microbiome of this imperiled amphibian species.

Our overarching objective was to characterize the gut microbial diversity and composition of eastern hellbenders in zoos and the wild. We investigated the bacterial gut assemblages of individuals pre-release (i.e., at the Nashville Zoo) and post-release (i.e., in native habitat), to characterize the temporal dynamics of gut microbial communities. We also characterized the gut microbiome of wild resident hellbenders (i.e., individuals that had never been held in zoos). Comparing wild residents with post-release individuals allowed us to evaluate the potential for microbiome restructuring after reintroduction. In addition, we tested if the introduction of a wild diet in headstart programs can contribute to rewilding of the microbiome prior to release. Lastly, we present the first characterization of the hellbender gut mycobiome as an initial attempt to understand how fungal taxa vary between hellbenders under human care and in wild settings.

We hypothesized that: 1) hellbender gut bacterial communities are structured by the animal’s environmental setting (zoo vs. wild), 2) the bacterial gut microbiome of hellbenders is altered by dietary rewilding, 3) upon introduction to the wild, the hellbender gut microbiome will shift over time to resemble wild individuals, and 4) the taxonomic composition of the mycobiome will be influenced by environmental setting. This study establishes a baseline knowledge of the previously uncharacterized gut microbiome of the imperiled eastern hellbender and identifies some of the complex dynamics that govern the gut microbiome in headstart populations at zoos, post-release, and in the wild, resulting in direct implications for conservation approaches.

## METHODS

### Study Sites – Zoos

The Nashville Zoo at Grassmere began their eastern hellbender conservation efforts in 2007, including a headstart program that began in 2015. Since 2015, zoo staff, along with university and state partners, have collected eastern hellbender eggs from streams in Middle Tennessee and reared them under human care until individuals were large enough for radio transmitter implants. Individuals were then released into the wild to assist with augmentation of current declining and genetically distinct populations of hellbenders in Middle Tennessee, and they were monitored with the goal of understanding the survival and spatial ecology of hellbenders in the geographic region. Zoo-reared individuals were housed in glass aquaria with carbon filtered municipal tap water with sodium thiosulfate (dechlorinator) added for make-up water. The filtration systems consisted of filter socks for mechanical filtration in addition to bio ball media for biological filtration. Two large racking systems received the same shared filtration and water and were made up of ∼20 aquaria. Water filtration and aquarium rack systems ranged from 120-400 total gallons capacity with river water collected from potential release sites being opportunistically added to the systems. Larvae and young juveniles were raised in 10-15 gallon glass or acrylic aquariums with multiple individuals (n ∼10-20) per aquarium. Larger juveniles and subadults/adults were housed in 20 gallon aquariums at lower densities (n ∼1-5). The density of animals in each aquarium was based on animal size, behavior, aquarium size, and space limitations. Each subadult and adult tank contained a gravel substrate along with half-cut PVC pipes as cover objects. Subadult and adult hellbenders were typically fed a zoo diet that primarily consisted of minnows, or other live fish, and thawed lake smelt, supplemented with nightcrawlers and shrimp.

In preparation for reintroduction into the wild, zoo staff gradually exposed hellbenders to river water from potential release sites via partial (∼30%) water changes, and the diets of individuals were also opportunistically supplemented with crayfish to serve as natural prey items. Tennessee Wildlife Resource Agency (TWRA) staff collected and transported crayfish and stream water from the watershed where the hellbenders were going to be released to minimize the potential for the introduction of novel pathogens. Additionally, a subset of individuals received an experimental antifungal terbinafine implant (see [46] for implant specifications) shortly prior to release as a form of prophylactic treatment with the intention of mitigating chytridiomycosis and other skin-associated fungal infections in the wild. Lastly, all pre-release hellbenders were implanted with series F1170 ATS model radio transmitters (Advanced Telemetry Systems, Isanti, MN, USA) and model APT12 passive integrated transponder (PIT) tags (Biomark, Boise, ID, USA) to aid in tracking animals post-release.

The other zoo population characterized in this study was from the Chattanooga Zoo. Most of the adult hellbenders were wild-caught as subadults/adults and maintained as permanent residents. These hellbenders were housed alone or co-housed with six other individuals in a large tank with filtered tap water, and they received a diet including but not limited to feeder fish and macroinvertebrates. As the Chattanooga Zoo hellbenders sampled in this study were not expected to be released as part of the headstart program, this population received no further environmental or dietary manipulations.

### Study Sites – Field

In Tennessee, eastern hellbenders are restricted to the eastern two-thirds of the state, namely within the Cumberland and Tennessee River drainages [41]. Thus, field sampling of wild and Nashville Zoo-released individuals only occurred at stream sites in portions of Middle and East Tennessee in the Interior Plateau and Blue Ridge ecoregions (Environmental Protection Agency level III), respectively. In Middle Tennessee, wild resident hellbenders and reintroduced individuals from the Nashville Zoo were sampled from release sites in the Lower Duck River watershed. In East Tennessee, wild hellbenders were sampled from a stream in the Cherokee National Forest.

### Sample Collection – Zoos

Individual hellbenders were removed from their respective tanks using nitrile gloves and placed in clean, sanitized plastic containers. Each individual was then identified via visual inspection (i.e., skin patterning) and PIT tag number. Hellbenders were then rinsed with ∼50 mL of 2-hr autoclaved water to remove any transient microbes and were subsequently wrapped in a clean, dampened microfiber towel to restrain the animal. Gut microbiome samples were collected from adult hellbenders by swabbing the cloaca using sterile rayon tipped swabs (Puritan Medical Products, Guilford, ME, USA, Ref # 25-806 1PR). Prior to swabbing, each swab was dipped in autoclaved mineral oil for ease of sample collection and to minimize any potential harm to the animal. Then, the swab head was rotated five times within the cloaca. Swabs were stored in an empty sterile microcentrifuge tube, immediately placed on ice, and later transferred to -80 °C storage upon return to Middle Tennessee State University (MTSU; Murfreesboro, TN). Any fecal samples were collected opportunistically with a sterile scoopula or siphoned out of the tank with a serological pipet and then transferred to a sterile microcentrifuge tube filled with 1 mL of sterile water. Morphometric data, including body mass (g) and total length (TL; cm), were also recorded for each hellbender following microbiome sampling. For Nashville Zoo hellbenders, sex was also recorded for each animal following a PCR-based assay outlined in [47]. To avoid any sample cross-contamination, nitrile gloves, plastic containers, and restraining towels were changed after handling each hellbender at both zoos. Hellbenders at the Nashville Zoo were sampled multiple times between the timeframe of February 2023 and April 2024, whereas sampling only occurred once at the Chattanooga Zoo in April 2024.

### Sample Collection – Field

Field sampling occurred during summer 2024 for Middle and East Tennessee. Headstart hellbenders released from Nashville Zoo were tracked at field sites in Middle Tennessee via radio telemetry. In addition to radio telemetry, we used a Biomark HPR Plus Reader equipped with an external antenna as an additional tracking method to verify and identify an individual hellbender’s relative location. Rock-lifting and snorkeling surveys were performed to capture zoo-released individuals once their exact location was determined. Additionally, we opportunistically surveyed for wild resident hellbenders with rock-lifting and snorkel survey techniques and used log peaveys to temporarily lift suitable cover rocks. All captured individuals were handled with a clean pair of field gloves, and upon capture, each hellbender was placed in a clean mesh bag before being transferred to a sanitized plastic container. Identification of zoo-raised animals was confirmed via PIT tag, and captured wild residents were implanted with a PIT tag for future population monitoring. In East Tennessee, wild resident hellbenders were captured during opportunistic surveys. Collection of gut microbiome samples followed the protocol outlined above with the addition that each hellbender’s cloaca was also rinsed with autoclaved water for an additional 15-20 seconds to remove any transient environmental microbes prior to sample collection. Only individuals greater than 20 cm TL were sampled in the field due to limitations of the size of the swab head relative to cloaca size.

### DNA Extraction and Amplicon Sequencing

DNA was extracted from a total of 197 cloacal swab samples (Nashville Zoo: n = 162; Chattanooga Zoo: n = 12; Middle TN Wild: n = 3; Middle TN Zoo Recapture: n = 4; East TN: n = 16) and 7 opportunistic fecal samples (Nashville Zoo: n = 3; Middle TN Wild: n = 2; Middle TN Zoo Recapture: n = 2) per the manufacturer’s protocol of the Qiagen DNeasy PowerSoil Pro kit. Negative control samples with no DNA template, consisting of sterile rayon tipped swabs dipped in mineral oil, were included in each round of DNA extraction to control for cross-contamination. For amplicon sequencing, the bacterial V4 region of the 16S rRNA gene was amplified for all samples (n = 197 cloacal swabs; n = 7 fecal samples) using the 515F and 806R primers [48] under the following thermal cycling conditions: initial denaturation at 98 °C for 2 min, followed by 35 cycles of denaturation at 98 °C for 10 s, annealing at 55 °C for 15 s, extension at 72 °C for 5 s, and final extension at 72 °C for 5 min. The fungal ITS1 rDNA gene was amplified using the ITS1FI2 and ITS2 primers [49, 50] with the following conditions: 98 °C for 2 min, followed by 35 cycles at 98 °C for 30 s, 52 °C for 30 s, 68 °C for 30 s, and 68 °C for 10 min. Only fecal samples (n = 7) were sequenced for the ITS1 gene due to issues with fungal amplification from cloacal swabs. The resulting amplicons were dual-indexed using the Nextera XT kit (Illumina, San Diego, CA, USA) according to the protocol outlined in [51]. Amplicons underwent size selection using MagBio HighPrep PCR magnetic beads (Gaithersburg, MD, USA), and gel electrophoresis was performed to evaluate amplicon size. The concentration of each DNA sample in the library was quantified using a Quantus fluorometer (Promega, Madison, WI, USA) to allow for normalization prior to sequencing. High-throughput amplicon sequencing was performed on an Illumina MiSeq platform (Illumina Inc., San Diego, CA, USA) using 3 total runs with 2 x 250 bp paired-end read chemistry.

### Bioinformatics

We generated a total of 16,119,530 raw bacterial sequence reads from amplicon sequencing. We conducted bioinformatic analysis of the bacterial communities of all samples using the mothur v1.48.0 software platform [52] according to the MiSeq Standard Operating Procedure (SOP) [53], with the modification that primer sequences were trimmed from reads after the formation of contigs, allowing up to one nucleotide difference in the forward primer sequence and up to three nucleotide differences in the reverse primer sequence. Sequences were aligned to the SILVA v138 reference database [54] for taxonomic assignment, and any unidentified sequences and those that were identified as chimeras or non-bacterial taxa were discarded prior to *de novo* clustering into operational taxonomic units (OTUs) according to 97% sequence similarity as a proxy for bacterial taxa. Any OTUs identified in no DNA template controls were removed as contaminants based on a 0.45 probability threshold using *decontam* [55]. One OTU (OTU00001; *Escherichia-Shigella*) was a suspected contaminant due to its prevalence across all samples, including controls, and had to be manually removed from the dataset prior to downstream analyses. This resulted in a data set with a total of 425,666 bacterial sequence reads retained after complete processing. Samples were rarefied to a depth of 6,000 reads (88% of total samples retained) using the *rrarefy* function in *vegan*. This final dataset was then used for downstream statistical analyses via R Statistical Software v4.4.1 [56].

Bioinformatics processing of the ITS amplicon sequences (298,031 total reads) generated from hellbender fecal samples was performed via the QIIME 2 v2023.9 software platform [57]. Demultiplexed paired-end sequences were trimmed to the ITS1 region using ITSxpress [58]. Sequences were then denoised and quality filtered to remove non-target reads and potential chimeric sequences using the DADA2 plugin [59]. The VSEARCH plugin [60] was utilized to perform *de novo* clustering of sequences into OTUs at 97% sequence similarity, and the sklearn classifier was applied to assign taxonomy to sequences according to the UNITE v9.0 database [61]. After quality filtering and taxonomic assignment, the fecal samples were subsampled at a depth of 21,000 sequence reads per sample, resulting in a total of 170,466 fungal sequences retained in the final dataset.

### Statistical Analysis

#### Zoo vs. Wild Microbiomes

Cloacal swab samples from zoo (Nashville Zoo: n = 137; Chattanooga Zoo: n = 12) and wild-caught hellbenders (Middle TN Wild: n = 3; Middle TN Zoo Recapture: n = 4; East TN: n = 16) were analyzed to investigate the impact of environmental setting (zoo vs. wild) on gut bacterial assemblages. As a note, six cloacal swabs from the Nashville Zoo were stored in 40% glycerol and used in these analyses as replacement samples due to sample type troubleshooting during molecular wet lab work. No significant differences in either alpha or beta diversity were detected between these glycerol stored samples and dry stored cloacal swabs (paired t-test, t = - 0.353, df = 5, p > 0.05) (Table S1). To measure differences in alpha diversity between all zoo and wild samples, we calculated bacterial OTU richness as Hill numbers (D; q = 0) using the *hill_taxa* function available in *hillR* [62] to represent the effective number of OTUs (i.e., a count of unique OTUs for bacterial richness in this case) as proposed by Jost [63]. To evaluate whether an animal’s environment affects taxonomic diversity, OTU richness was compared between zoo and wild-caught hellbenders using a generalized linear mixed model (GLMM) approach via *glmmTMB* [64]. Corrected Akaike Information Criterion (AICc) values were used to account for small sample sizes and inform model selection, in which models with the lowest AICc value were favored. Scaled residuals simulated using *DHARMa* [65] were visualized to evaluate model fit and check model assumptions of the AICc-selected model. In the final GLMM, a negative binomial distribution with quadratic parameterization and log link was used to manage data overdispersion. The environmental setting of each individual during sample collection (i.e., zoo or wild) and total body length (TL) were included as fixed effects, whereas random effects consisted of sample collection date and animal ID. The marginal coefficient of determination (R^2^) of the model was calculated using the *r.squaredGLMM* function within *MuMIn* [66] to ascertain how much of the model variance was explained by the fixed effects.

To visualize the shared and unique bacterial families among zoos and wild settings, a network was constructed and visualized with Cytoscape v3.10.3 [67]. Family level absolute abundance counts were used to generate edges and nodes for Cytoscape input files using a custom R script. Family core microbiome heatmaps were generated using the *plot_core_microbiome_custom* function with default settings (prevalence values ranging from 0.01 to 1, detection threshold ranging from 0.001 to 0.03, and a minimum prevalence value cutoff of 20% across samples) from the R package *DspikeIn* [68].

Bacterial community structure between zoo and wild-caught hellbenders was visualized using non-metric multidimensional scaling (NMDS) for both Bray-Curtis and Raup-Crick dissimilarity metrics using *vegan* [69]. The full dataset was randomly subsampled to include one sample at a randomly selected sampling date for every hellbender sampled at the Nashville Zoo (n = 50) to account for potential pseudoreplication in community structure since Nashville Zoo individuals were repeatedly sampled. To evaluate if there were significant differences in multivariate dispersion between the zoo population and wild groups, a permutational test of multivariate homogeneity of groups dispersions was performed for both dissimilarity metrics using the *betadisper* function in *vegan* [69]. If significant differences were detected, a Tukey’s Honest Significant Difference test was conducted to evaluate any significant pairwise group comparisons. If multivariate dispersion was significantly greater for groups with a larger sample size, permutational multivariate analysis of variance (PERMANOVA) tests proceeded as planned. However, if greater dispersion was present for smaller groups, random subsampling for equal sample sizes was performed to create a balanced dataset, as PERMANOVA is robust to heterogeneity of multivariate dispersion for balanced sampling designs [70]. The *adonis2* function was used to perform a PERMANOVA with 999 permutations for both dissimilarity metrics to assess whether average community structure differed between zoo and wild hellbenders. Due to the inclusion of multiple sites in the analysis, spatial autocorrelation in the bacterial assemblages was accounted for by stratifying permutations according to sample collection site. The interaction between bacterial richness and an animal’s broad geographic location (i.e., the animal’s zoo location or region of Tennessee, with recaptured and wild residents considered separately) as well as the main effect of the animal’s environmental setting (zoo vs. wild) were included in both the Bray-Curtis and Raup-Crick PERMANOVAs.

#### Effects of a Diet Manipulation at the Nashville Zoo

Cloacal swab samples collected from males within the same cohort that hatched in 2018 at the Nashville Zoo were analyzed based on dietary treatments. Only male animals were sampled to control for effects of sex, age, and potential genetic contributions on the gut microbiome, as the eggs for individuals within a cohort are typically collected at the same field site or under the same nest rock. Based on differential crayfish feeding across three sampling time points (May 2023, November 2023, and January 2024), samples were divided into two groups (Group 1: n = 28; Group 2: n = 26), where the diet of Group 1 was supplemented with crayfish (e.g., a wild diet) for the last two sampling points, whereas individuals in Group 2 had only been fed crayfish by the last time point.

A GLMM with a Poisson distribution was used to investigate the effect of a wild diet on gut bacterial richness. Prior to modeling the data, scaled mass index (SMI) was calculated from individual body mass and TL according to [71], which serves as an important proxy for host body condition and health in hellbenders [72]. Fixed effects of the model included the main effects of SMI, sample collection date, feeding group, and the interaction between sample collection date and feeding group. Random effects consisted of animal ID and the date of DNA extraction, both of which were included as random intercepts in the model. Model selection and evaluation of model fit followed the steps explained previously.

Bacterial community structure, as measured by Bray-Curtis and Raup-Crick dissimilarities, for both male feeding groups at each sampling time point was visualized and statistically analyzed via NMDS plots and PERMANOVAs, respectively. The date of sample collection and the interaction between bacterial richness and feeding group were assessed as predictors for the Bray-Curtis PERMANOVA, whereas feeding group was the only predictor used for the Raup-Crick PERMANOVA. Bacterial richness and date of sample collection were not evaluated due to poor model fit in the Raup-Crick matrix.

To further evaluate patterns in community structure that were observed upon introduction of a wild diet, individuals from both feeding groups were analyzed at the first and third sampling time point (May 2023: n = 14; Jan. 2024: n = 21) to serve as a reference before and after crayfish feeding. To determine the mechanistic processes governing beta diversity within this framework, total beta diversity was partitioned into turnover and nestedness [73, 74]. Values for turnover (β_SIM_), nestedness (β_SNE_), and total beta diversity (β_SOR_) were calculated with the Sørensen index using the *betapart* package and plotted as a density distribution plot as described in [75]. Briefly, the sample by OTU matrices were transformed to presence/absence data prior to being converted to a betapart object using the *betapart.core* function. A total of 12 before and 12 after crayfish feeding samples were randomly selected and resampled 100 times using the *beta.sample* function to generate the distributions for the three beta diversity measures. Twelve samples were chosen for this analysis to allow for random selection of samples within the before crayfish feeding dataset (May 2023; n = 14), which had a smaller sample size than the after crayfish feeding dataset.

To elucidate if particular bacterial taxa significantly shifted in relative abundance following the introduction of a natural prey item in a zoo setting, a Linear discriminant analysis Effect Size (LEfSe) analysis was performed for pre- and post-crayfish feeding. The LEfSe analysis was conducted using the *microbiomeMarker* package [76] to evaluate differentially abundant/enriched taxa present in the gut microbiome before and after wild diet supplementation.

#### Restructuring of the Microbiome Post-Release

Sample size of recaptured hellbenders from the Nashville Zoo (n = 4) was limited due to challenges in field capture of released animals as well as due to the fact that repeated disturbance of headstarted animals can adversely impact survival after translocation. Therefore, statistical analyses were restricted. The relative abundances of bacterial genera of post-release samples were qualitatively compared to pre-release samples from only two individuals with paired samples. We also included wild resident hellbenders from both Middle (n = 3) and East Tennessee (n = 16). These comparisons enabled us to identify how particular gut bacteria change from zoos to the wild and if post-release hellbenders share bacteria with wild residents.

#### Gut Mycobiome

Opportunistically collected fecal samples (Nashville Zoo: n = 3; Middle TN Wild: n = 2; Middle TN Zoo Recapture: n = 2) were used to evaluate the taxonomic diversity of fungi present within hellbender gut assemblages. Due to the small sample size, we were limited to a descriptive exploration of the relative abundance of fungal genera across all individuals, which was coupled with a preliminary investigation of fungal diversity using Hill numbers.

## RESULTS

### Zoo vs. Wild Microbiomes

After quality filtering, ∼2,200 bacterial OTUs were identified from zoo and wild hellbender cloacal swabs, representing 29 phyla, 62 classes, and 159 orders. The most dominant phyla in the hellbender gut microbiome were Proteobacteria (46.9% relative abundance), Bacteroidota (33.4%), Firmicutes (10.3%), and Actinobacteriota (4.7%) while all other phyla represented less than 3%. However, at a finer taxonomic scale, the composition of the gut microbiome appeared to vary according to bacterial order between zoo and wild-caught populations (Fig. 1).

**Figure 1.**
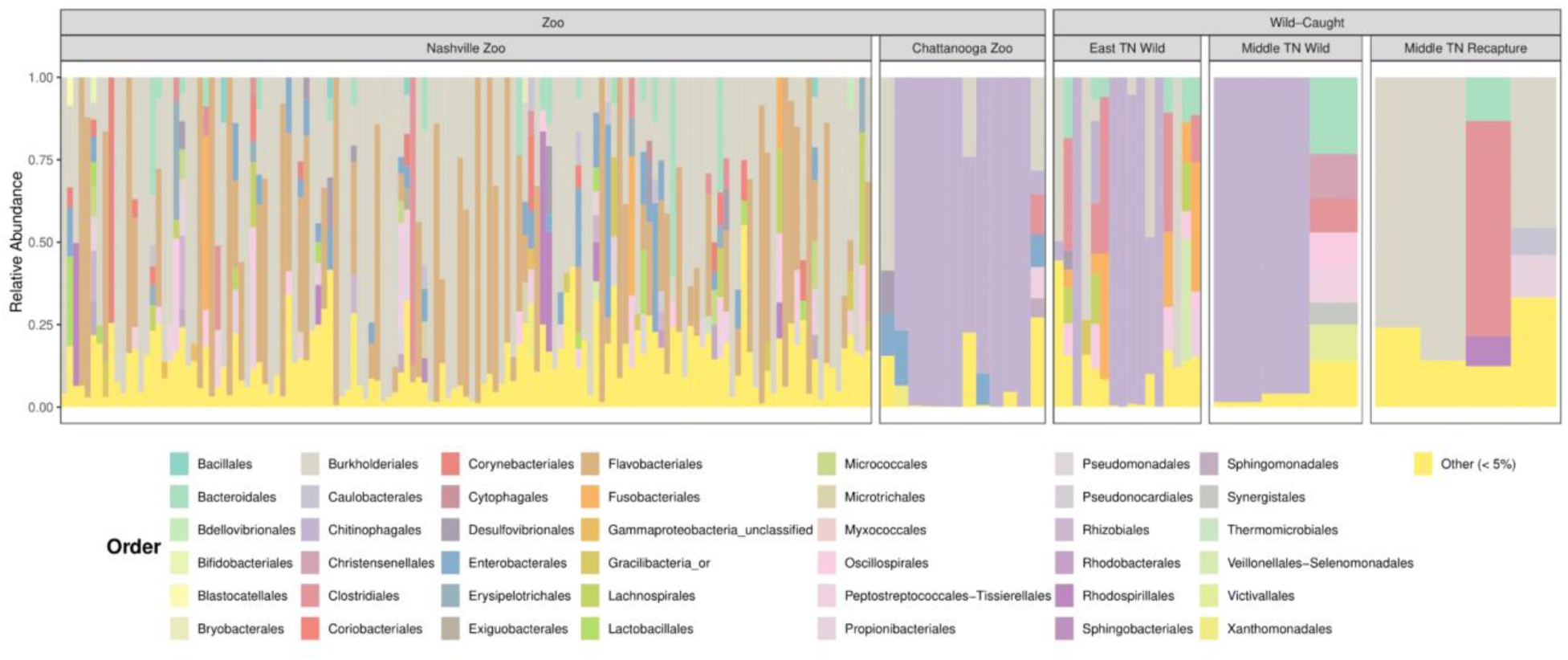
Taxonomic composition of zoo and wild-caught hellbender gut assemblages. Relative abundance of bacterial orders across zoo and wild settings. Each bar represents an individual sample, and taxa representing less than 5% relative abundance were classified as Other.

Nashville Zoo hellbenders were dominated by the Burkholderiales (43.2%) and Flavobacteriales (24.6%) orders, while Chattanooga Zoo hellbenders were dominated by Chitinophagales (76.8%). Chitinophagales was also present in high abundance in wild hellbenders from East (41.9%) and Middle Tennessee (65.0%), whereas Burkholderiales (17.0%) was the second most abundant order in East Tennessee hellbenders. Post-release hellbenders were largely composed of Burkholderiales (51.8%) and Clostridiales (16.7%).

Network analysis displayed the partitioning of family level OTUs across zoo and wild-caught individuals (Fig. 2a). Of the 407 different bacterial families detected, only 185 were shared between zoo and wild individuals, whereas 163 were unique to zoo individuals and 59 to wild individuals. Several families showed remarkable abundance shifts between groups. For example, Weeksellaceae, Eggerthellaceae, and Chromobacteriaceae were more abundant in the zoo group, whereas Barnesiellaceae, Selenomonadaceae, and Synergistaceae were enriched in the wild group.

After defining the core microbiome, significant differences were observed between zoo and wild hellbenders (Fig. 2b). The core microbiome of zoo individuals comprised 18 bacterial families in total, with Oxalobacteraceae, Weeksellaceae, Propionibacteriaceae, and Chitinophagaceae identified as prevalent and/or abundant families (Fig. 2b). In contrast, the core microbiome of wild-caught hellbenders was more diverse and more consistently shared, with 44 families present in at least 20% of the samples. Notably, members of Chitinophagaceae appeared to be present at relatively greater prevalence and abundance in wild hellbenders (Fig. 2b).

**Figure 2.**
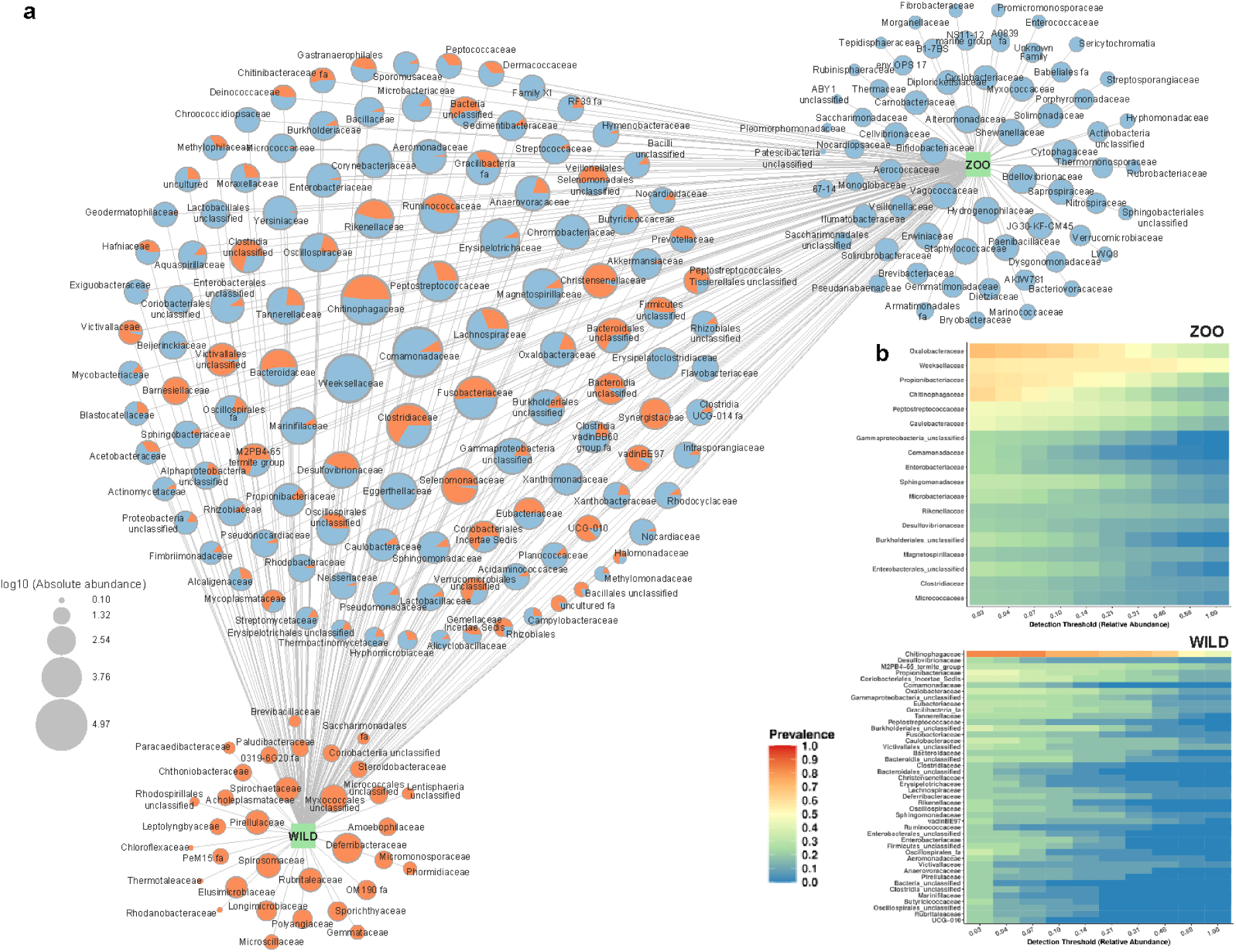
Family-level network and hellbender core bacterial microbiome. a) Shared and unique bacterial families between zoo and wild individuals. b) Bacterial families identified as core gut microbiome members in zoo and wild individuals based on both prevalence and relative abundance.

In addition to differences in taxonomic composition, individuals in zoos featured significantly lower bacterial richness than wild-caught individuals (GLMM, z-value = -2.597, p < 0.01), while TL did not significantly impact bacterial richness (GLMM, z-value = 0.691, p > 0.05) (Table S2).

For Bray-Curtis distances, group dispersion was similar between zoo and wild individuals (permutest, F_1,83_ = 0.929, p > 0.05) but was significantly different across broad geographic location (permutest, F_4,80_ = 7.591, p < 0.001), with Chattanooga Zoo hellbenders having less dispersion than Nashville Zoo and wild East TN hellbenders. Neither environmental setting (permutest, F_1,83_ = 0.001, p > 0.05) nor broad geographic location (permutest, F_4,80_ = 2.112, p > 0.05) significantly influenced Raup-Crick group dispersion. For the Bray-Curtis metric (Fig. 3a), bacterial community structure was influenced by environmental setting (PERMANOVA, F_1,75_ = 3.766, R^2^ = 0.039, p < 0.01) and the interaction between richness and broad geographic location (PERMANOVA, F_4,75_ = 2.061, R^2^ = 0.085, p < 0.001) but not by the main effects of bacterial richness (PERMANOVA, F_1,75_ = 1.019, R^2^ = 0.011, p > 0.05) or broad geographic location (PERMANOVA, F_3,75_ = 2.925, R^2^ = 0.091, p > 0.05). However, the Raup-Crick metric indicated that environmental setting (PERMANOVA, F_1,75_ = 3.326, R^2^ = 0.024, p < 0.01), broad geographic location (PERMANOVA, F_3,75_ = 9.564, R^2^ = 0.206, p < 0.05), and the interaction between bacterial richness and broad geographic location (PERMANOVA, F_4,75_ = 7.032, R^2^ = 0.202, p < 0.01) influenced community structure while richness did not (PERMANOVA, F_1,75_ = 4.305, R^2^ = 0.031, p > 0.05) (Fig. 3b).

**Figure 3.**
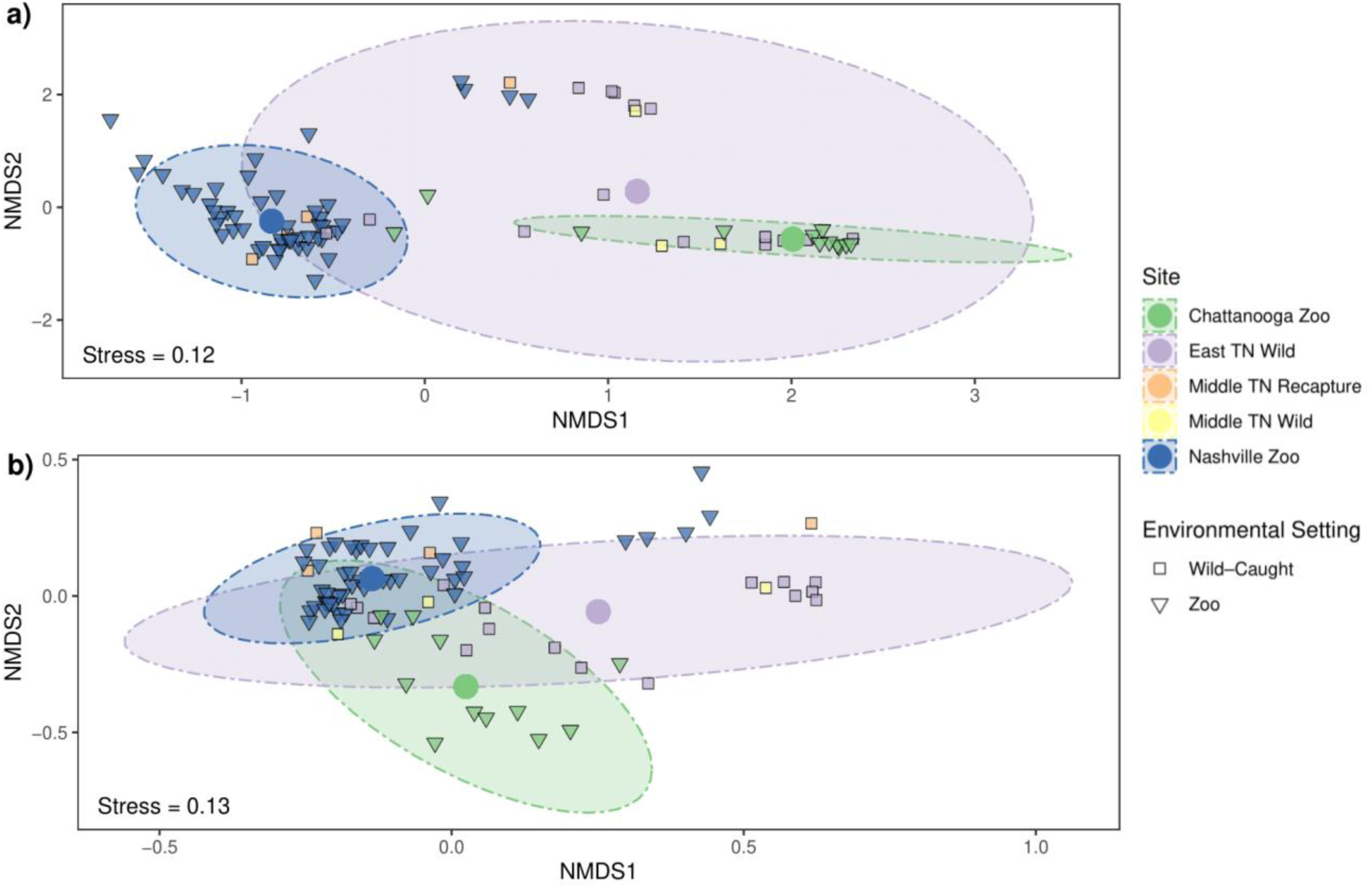
Bacterial community structure of zoo and wild hellbenders. Non-metric multidimensional scaling ordinations according to Bray-Curtis (a) and Raup-Crick (b) dissimilarity metrics. Due to repeated sampling at Nashville Zoo, samples collected from Nashville Zoo were subsampled to include one sample per animal. Ellipses represent 95% confidence intervals, and group centroids are shown as colored dots to represent the average community structure for hellbenders collected from each relative site.

### Effects of a Diet Manipulation at the Nashville Zoo

For male hellbenders analyzed as part of the wild diet manipulation, bacterial richness significantly varied according to collection date (GLMM, z-value = -6.508, p < 0.001), feeding group (GLMM, z-value = -3.047, p < 0.01), SMI (GLMM, z-value = 2.305, p < 0.05), and the interaction between sample collection date and feeding group (GLMM, z-value = 3.599, p < 0.001) (Fig. S1; Table S2). For Bray-Curtis dissimilarity, some similar patterns were revealed, with collection date (PERMANOVA, F_2,48_ = 1.933, R^2^ = 0.067, p < 0.05), feeding group (PERMANOVA, F_1,48_ = 1.923, R^2^ = 0.034, p < 0.05), and the interaction between bacterial richness and feeding group (PERMANOVA, F_1,48_ = 2.02, R^2^ = 0.035, p < 0.05) being significant predictors of average community structure. However, richness was not a significant predictor of community structure (PERMANOVA, F_1,48_ = 1.496, R^2^ = 0.026, p > 0.05; Fig. 4a). While the homogeneity of variance assumption was not violated for feeding groups (permutest, F_1,52_ = 0.104, p > 0.05) or across collection dates (permutest, F_2,51_ = 0.410, p > 0.05) using Bray-Curtis dissimilarity, this assumption was violated for feeding group using Raup-Crick dissimilarity (permutest, F_1,52_ = 14.325, p < 0.001). Consequently, random subsampling of the dataset was performed to result in an equal number of individuals in each feeding group (n = 26). After subsampling, we detected significant differences in dispersion between feeding groups (permutest, F_1,50_ = 15.303, p < 0.001). Nevertheless, Raup-Crick dissimilarity showed that feeding group (PERMANOVA, F_1,50_ = 13.138, R^2^ = 0.208, p = 0.05) had a significant effect on the structure of bacterial communities (Fig. 4b).

**Figure 4.**
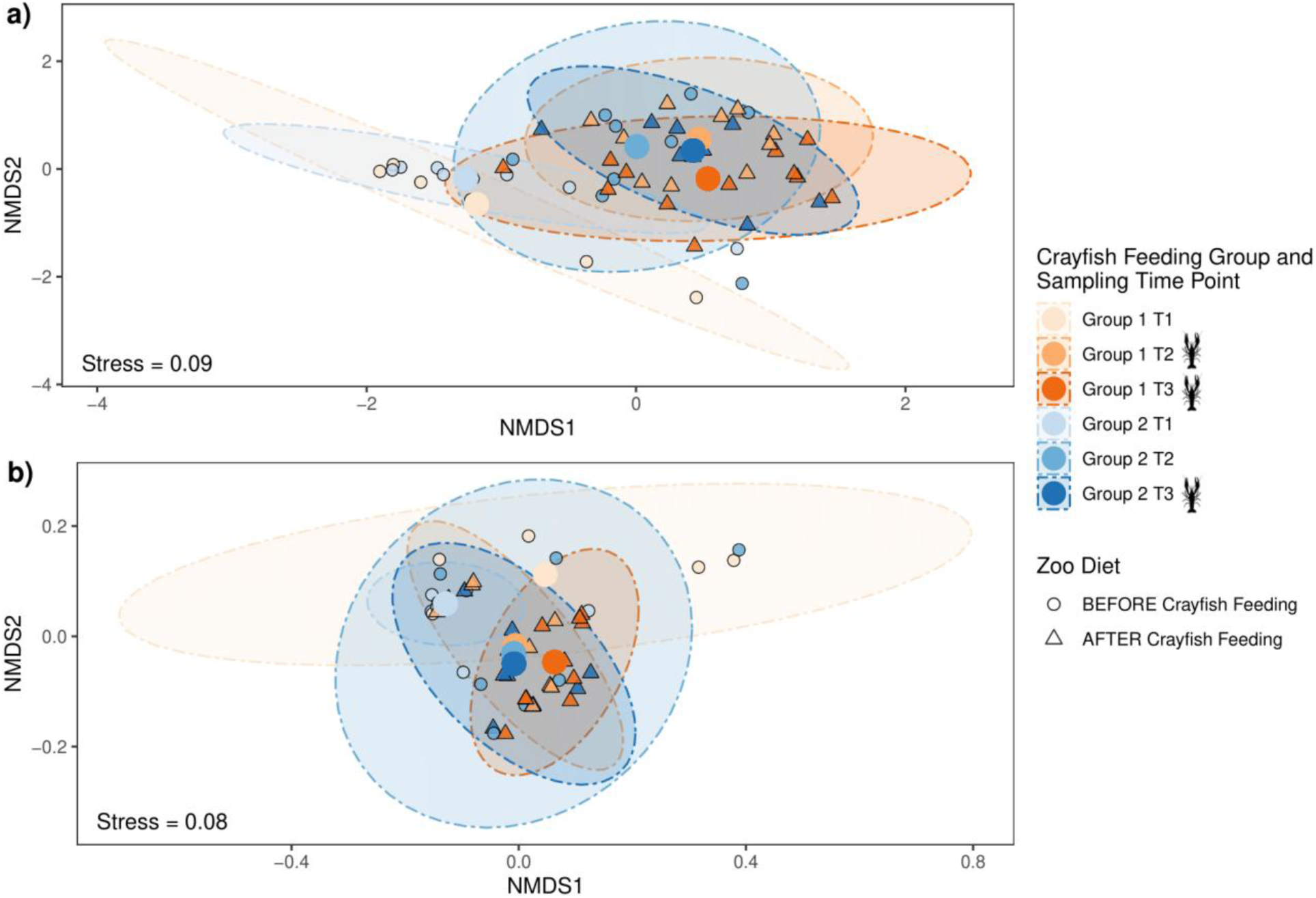
Community structure along a temporal gradient after the introduction of a natural prey item. Non-metric multidimensional scaling ordinations of bacterial communities of male hellbenders exposed to a wild diet (crayfish) at the Nashville Zoo using Bray-Curtis (a) and Raup-Crick (b) dissimilarity matrices. Ellipses represent 95% confidence intervals, and group centroids are shown as colored dots to represent the average community structure of the two crayfish feeding groups across different sampling dates (T1 = May 2023, T2 = November 2023, T3 = January 2024).

Examining the separate mechanisms that comprise beta diversity revealed that turnover and nestedness differed before and after introducing crayfish as part of a natural zoo diet (Fig. S2). Total beta diversity was comparable before (mean 0.904 ± 0.006 SD) and after (0.892 ± 0.004) crayfish feeding. Nestedness was relatively low throughout but was greater before (0.076 ± 0.013) crayfish feeding rather than after (0.023 ± 0.006). In contrast, turnover increased after crayfish feeding (0.869 ± 0.006) compared to before (0.828 ± 0.012).

Based on the LEfSe analysis, differentially abundant taxonomic features were broadly clustered phylogenetically according to crayfish feeding status (Fig. 5a). Of these identified features, a total of 11 bacterial genera were significantly enriched before crayfish feeding whereas 10 genera were enriched after crayfish feeding (Fig. 5b).

**Figure 5.**
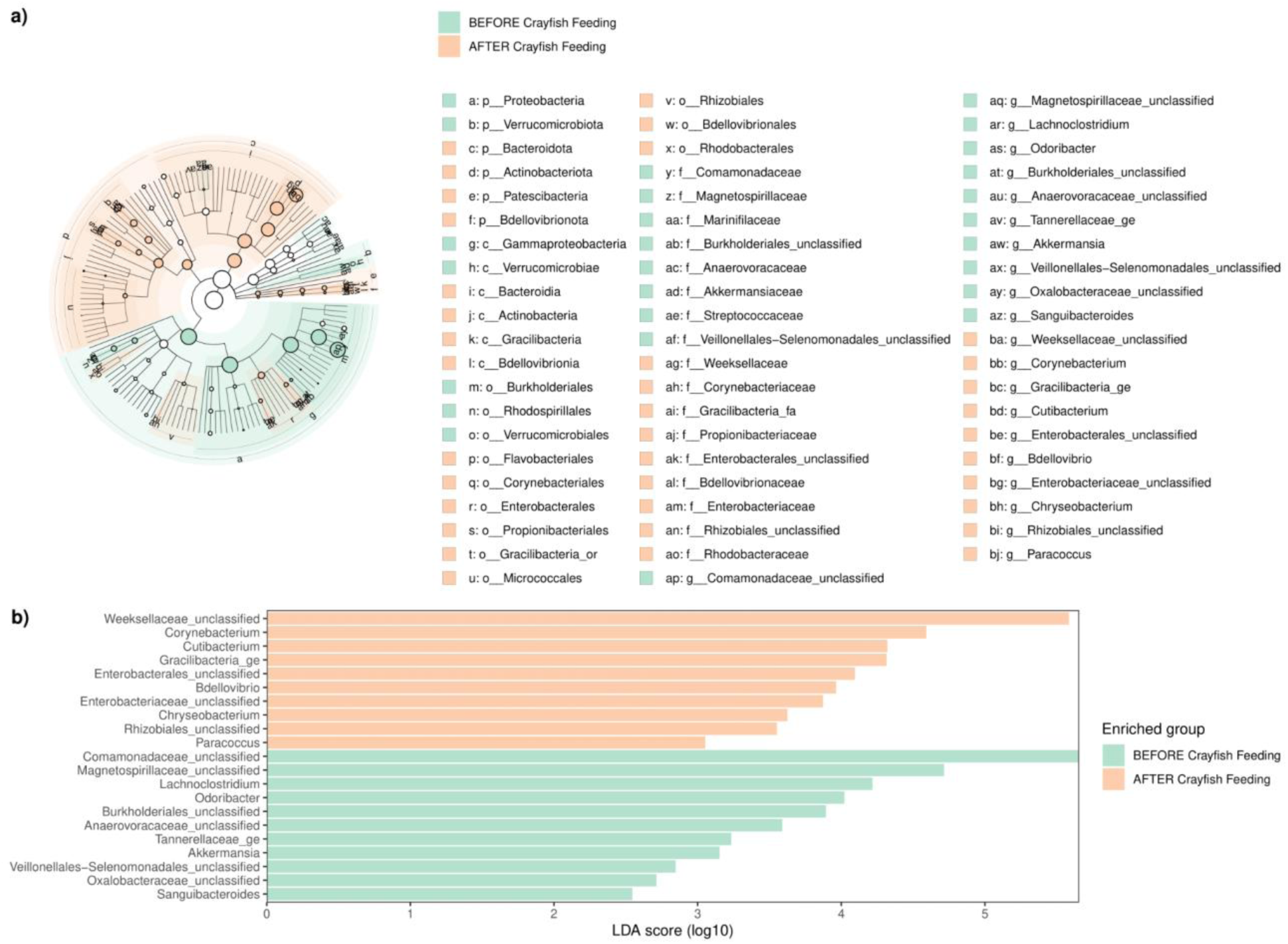
Differentially abundant bacterial taxa associated with crayfish feeding. Taxa enriched in the gut microbiome of male hellbenders at the zoo before and after introduction of a wild diet (crayfish) according to linear discriminant effect size analysis. a) Cladogram of differentially abundant taxonomic features. b) Histogram of enriched bacterial genera.

### Restructuring of the Microbiome Post-Release

The bacterial composition of hellbenders reared at Nashville Zoo shifted following introduction to the wild (Fig. 6a). Prior to release, two hellbenders at the zoo primarily consisted of unclassified Weeksellaceae (23.5% of total relative abundance), *Vogesella* spp. (16.6%), and unclassified Comamonadaceae (9.4%). Upon recapture in the wild, all individuals were dominated by unclassified Comamonadaceae (49%) and unclassified Clostridiaceae (14.9%). These taxa ultimately varied from the most abundant bacteria in wild residents, in which unclassified Chitinophagaceae was the predominant taxon found in Middle (65.0%) and East Tennessee (41.8%) hellbenders. Additionally, unclassified Comamonadaceae and *Cetobacterium* spp. comprised 15.7% and 7.7% abundance, respectively, in East Tennessee individuals.

Post-release hellbenders shared a greater number of OTUs with wild residents than with pre-release samples at Nashville Zoo (Fig. 6b). Out of a total of 1,159 OTUs, only 37 were shared between pre-release and post-release samples. Conversely, post-release samples shared 98 and 210 OTUs with Middle and East Tennessee wild residents, respectively. Approximately 198 OTUs were unique to post-release hellbenders.

**Figure 6.**
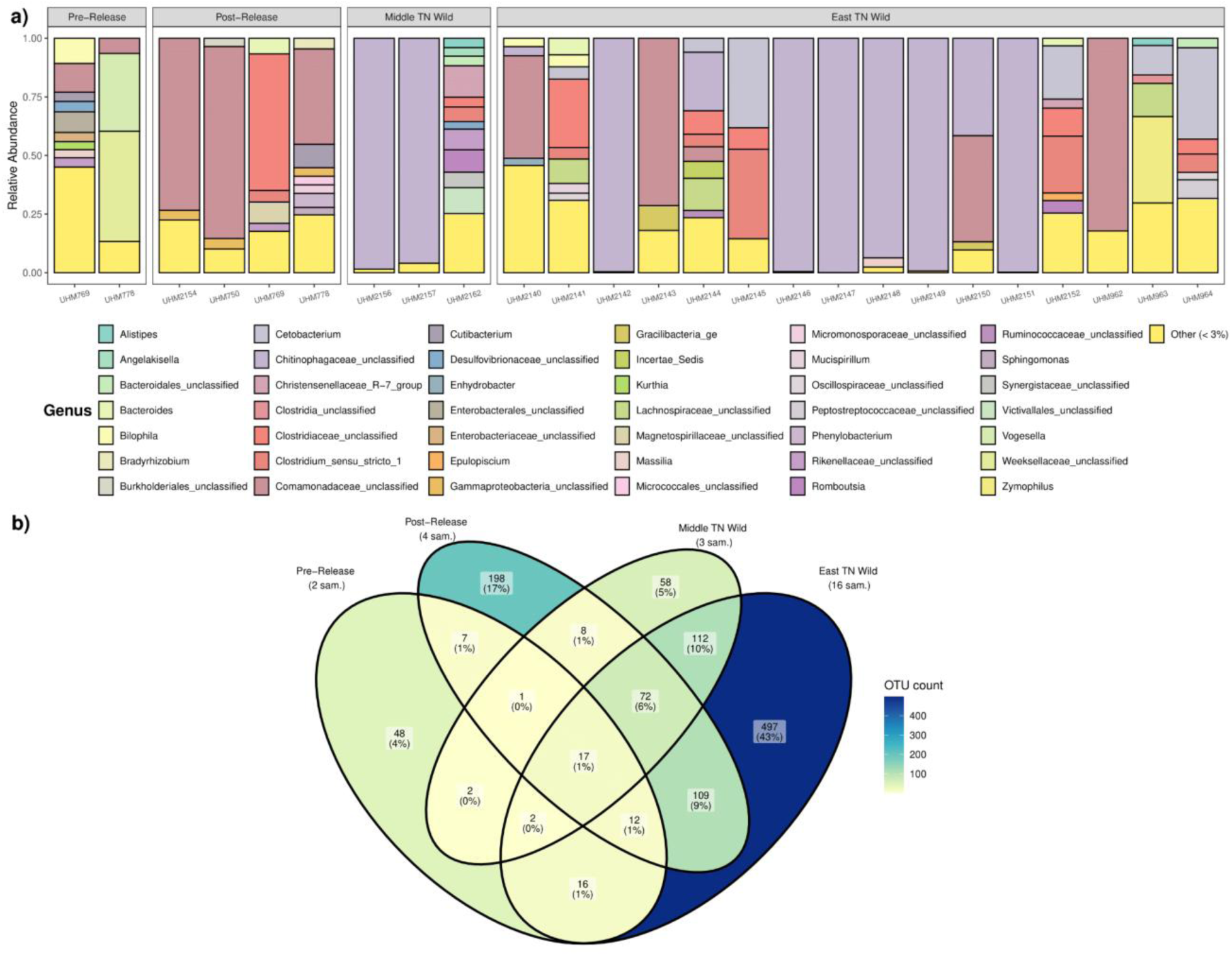
Assemblage composition of zoo-released recaptures and wild residents. a) Relative abundance of bacterial genera for Nashville Zoo-reared hellbenders pre-release (i.e., at the zoo) and post-release (i.e., recaptured in the wild) along with wild hellbender populations. Bars represent individual hellbenders. Individual taxa representing less than 3% relative abundance were grouped as Other. b) Venn diagram representing the total number of OTUs shared between pre- and post-release Nashville Zoo individuals and wild resident hellbenders.

### Gut Mycobiome

Approximately 700 OTUs represented the gut mycobiome of Nashville Zoo and Middle TN wild-caught hellbenders. However, only ∼200 OTUs could be assigned taxonomy for statistical analysis, which represented 9 phyla, 26 classes, and 49 orders. Average fungal richness and variability differed between zoo (mean 27 ± 4.6 SD) and wild-caught individuals (40.8 ± 32.3; Fig. S3a). Only 11 of 182 OTUs were shared between zoo and wild individuals (Fig. S3b). In contrast, recaptured samples shared 21 OTUs with wild residents, and a total of 14 OTUs were found in both zoo and recaptured hellbenders.

The relative abundance of fungal genera also differed between zoo and wild individuals (Fig. S4). In pre-release headstart hellbenders, *Basidiobolus* (73.4%) and an unclassified Mortierellaceae (9.5%) were the most abundant members of the mycobiome. Both recaptured and wild residents exhibited similar abundances of unclassified Rozellomycota (Middle TN Zoo Recapture: 39.5%; Middle TN Wild: 35.8%). *Rhodosporidiobolus* was also identified in both recaptures (5.7%) and wild Middle Tennessee hellbenders (18.3%), while *Mucor* (13.9%) was abundant in wild residents.

## DISCUSSION

Characterizing the gut microbial assemblage dynamics of imperiled species across zoo and wild settings, including after translocation into their natural habitat, can have implications for conservation practices [12, 77]. A zoo setting had a strong effect on the hellbender gut microbiome, leading to reduced bacterial and fungal richness along with a different taxonomic composition and bacterial community structure compared to wild-caught individuals. Introduction of a natural diet in zoo-reared hellbenders impacted the community structure as a result of shifts in bacterial richness. On release to native habitats, the composition and relative abundance of both bacterial and fungal taxa shifted, demonstrating that introduction into a wild environment altered zoo microbiomes and may lead to the acquisition of natural microbiota.

Both alpha and beta diversity of the hellbender gut microbiome was significantly affected by zoo husbandry. These results contribute to growing knowledge that conditions present in zoo and wild environments play a key role in gut microbiome dynamics, which has been demonstrated across mammals [25, 29, 32], reptiles [27, 28, 78], amphibians [37], and birds [26]. Significantly lower bacterial richness was observed in zoo individuals compared to wild-caught hellbenders, which is largely consistent with the microbiome literature [10, 23, 26, 78, 79]. In general, greater microbial diversity is typically considered a measure of enhanced host health and fitness, contributing to immunity, increasing metabolic functionality, and promoting microbiome stability [80–82], although this assumption has recently been contested [83]. Additionally, our results showed that bacterial communities significantly differed between zoo and wild-caught hellbenders when emphasizing microbial abundance (Bray-Curtis) and when accounting for differences in alpha diversity (Raup-Crick). This suggests that alpha diversity differences between zoo and wild individuals may not be a key driver of the differences in beta diversity. These results may influence zoo husbandry practices because such compositional changes of the microbiome care can have important, potentially adverse implications for host health and function [25, 30, 79, 84–86], and may negatively impact headstart programs or other conservation efforts.

Comparing the abundance of microbial taxa across various environments can reveal key insights about the gut microbiome, as abundant and rare taxa often vary in how they contribute to microbial community dynamics [87, 88]. Overall, the relative abundance of bacterial orders for Chattanooga Zoo hellbenders resembled that of wild individuals, with Chitinophagales being the most abundant taxonomic order. This could be important considering the family Chitinophagaceae was identified as a core taxon in the hellbender gut microbiome. The similar abundance of Chitinophagales between Chattanooga Zoo hellbenders and wild residents could either be related to husbandry practices or greater resiliency of a wild-type gut microbiome to changes associated with human rearing. Since most of the Chattanooga Zoo adults were wild residents at one point, the core, wild-derived microbiota could be retained after being relocated to a zoo setting, which has also been documented in other host species [17, 89]. However, a comparison between Chattanooga Zoo adults sampled for this study and individuals reared solely at the zoo would be needed to help support this hypothesis.

The family Chitinophagaceae was identified as part of the core microbiome, particularly in wild hellbenders. Core microbiomes represent microbial taxa that are commonly found across hosts and/or environments, and are suggested to play an important part in host health and function [90, 91]. Taxa in the Chitinophagaceae family are predominantly associated with the degradation of complex carbohydrates, namely chitin [92, 93], and have been detected in the gut microbiome of multiple freshwater and marine fish species [94–96]. Chitinophagales were also found to increase in relative abundance in aquaculture-raised European sea bass fed an experimental diet supplemented with commercial shrimp chitin [96]. Chitinophagaceae have also been associated with many aquatic macroinvertebrates [95, 97–99], including being identified as a dominant member of the crayfish gill microbiome [100]. Thus, the general chitinolytic activity of Chitinophagaceae, along with their close association with crayfish, signify that these bacteria may not only originate from the diet but also may play a key role in the digestion of crayfish, which is the primary prey item for hellbenders in natural environments [101].

In addition to evaluating the gut microbiome across environmental contexts, investigating the effects of commercial diets are also important for conservation initiatives since zoo diets may not be reflective of dietary items consumed in wild conditions [12, 13, 77]. This may be particularly important for organisms with complex or specialized diets, whose gut microbiomes are often more affected by the lack of natural diets while under human care [102–105]. In this study, introduction of a natural prey item in a headstart program altered both alpha and beta diversity along a temporal gradient. The zoo diet led to increased nestedness among individuals while the introduction of a natural prey item promoted turnover within bacterial communities. This suggests that zoo diets promote some degree of homogenization of gut microbial assemblages [106–108], yet a crayfish diet increased both losses and gains of microbial taxa. Compositional shifts in the gut microbiome associated with implementing a natural diet have also been observed in mammalian models [35, 103] and have been linked to the retention of natural microbiota while animals are in human care [109]. The microbial species replacement associated with a natural diet in this study may improve functional diversity and strengthen resilience and stability of the hellbender gut microbiome while increasing chances of survival upon release into the wild by promoting initial rewilding of the microbiome.

Several of the differentially abundant taxa that were associated with standardized headstart program diets may represent potential pathogens or indicators of microbiome disturbance. To illustrate, Oxalobacteraceae was enriched in hellbenders fed a zoo diet and was discovered to be a positive gut biomarker for biliary tract cancer in humans [110], while *Odoribacter* was identified as a possible opportunistic pathogen in the gut microbiota of alpine musk deer in a zoo [111]. In contrast, multiple bacterial genera associated with crayfish feeding in a hellbender headstart program may serve as beneficial members of the microbiome. For example, *Gracilibacteria* has been associated with healthy oral microbiomes in humans [112], whereas members of *Cutibacterium* have been identified as early gut colonizers of healthy human infants [113] and are capable of producing key metabolites, including short-chain fatty acids [113, 114], peptides [115], and vitamins [114] in vertebrate and invertebrate species. These results suggest that implementation of a wild diet in zoos may lead to improved taxonomic diversity and functionality of the microbiome.

Despite a limited sample size, we found that zoo-released recaptures shared more OTUs with wild residents than with their pre-release samples at the zoo. This suggests that the released hellbenders have acquired microbiota from the wild environment that were absent in the zoo. Greater similarity between wild and headstarted conspecifics released into the wild have also been documented for white-footed mice [35] and Tasmanian devils [32]. Since many of the shared OTUs between recaptures and wild-caught individuals were not evident in the relative abundance plot, they likely represent rare OTUs. Differences in the diversity of rare taxa between zoo and wild populations have been detected in other vertebrate hosts [116], which may be important to consider since rare taxa can play a key role in microbial community dynamics [117]. As the first study to investigate the gut microbiome of hellbenders during a reintroduction effort, these preliminary results can serve as a baseline for future studies and suggest some evidence of restructuring of the hellbender gut microbiome post-release. However, more extensive field sampling of zoo-released individuals would be necessary to fully support these findings, which may also prove challenging with the consideration that field recapture may negatively impact survival by increasing stress on the animal during the capture process and by affecting long-term cover rock/site fidelity.

In addition to the bacterial microbiome, the host mycobiome can also be predictive of overall health [118–120], though fungal taxa remain relatively understudied in the gut of vertebrate hosts [119, 121, 122]. This particular knowledge gap is also demonstrated in our study’s results, in which approximately 500 fungal OTUs were unable to be taxonomically classified and could potentially represent an unexplored reservoir of fungal diversity. Similar to the bacterial microbiome, the mycobiome of wild-caught individuals exhibited greater richness than zoo individuals, which has been documented in other wildlife [123, 124]. Also, more fungal taxa were shared between recaptured hellbenders and wild residents, showing potential evidence of restructuring of the mycobiome upon release. The genus *Basidiobolus* was among the most abundant taxa in the hellbender mycobiome. *Basidiobolus* is a filamentous fungus that has previously been identified as a ubiquitous member of various reptile and amphibian gut assemblages [125–128] and acts as a keystone species by promoting microbiome stability [68] and shaping bacterial community dynamics [127]. *Basidiobolus* was enriched in the gut of zoo-reared hellbenders which may be reflective of certain diet items in ex-situ conservation populations, particularly macroinvertebrates, since the fungus is known to closely associate with arthropods during its life cycle [129]. Further research will be essential in comprehensively characterizing the hellbender gut mycobiome and drawing more reliable conclusions, but the results presented here can serve as a base for future work to establish an understanding of hellbender gut fungal diversity.

## CONCLUSIONS

This study provides the first characterization of the gut microbiome of eastern hellbenders, using both zoo-raised and wild-caught individuals. Though headstaring hellbenders in a zoo appears to impose a strong effect on the hellbender gut microbiome, implementing a wild diet alters gut microbial communities and may create conditions that both potentially improve host health and increase chances of survival upon reintroduction. However, additional rewilding strategies may be needed to promote more complete restructuring to a wild-type gut microbiome in headstart programs and ultimately requires long-term monitoring to confirm positive effects on host health. Despite sample size limitations, both the gut bacterial and fungal communities of zoo-raised individuals demonstrated the capacity to shift following reintroduction into native habitat, exhibiting sets of natural microbiota that were shared with wild residents. Future work is needed to ultimately elucidate and connect the microbiome changes identified in this study to host health and individual survivability in the wild. Nonetheless, the results of this study may act as a foundation to advise current husbandry management practices and advance the long-term conservation of this charismatic salamander.

## Acknowledgements

The authors would like to specially thank Kylie Moe for providing training on molecular wet lab techniques. We would also like to extend a sincere thank you to Kaitlyn Murphy for providing general project feedback and assistance with sample collection at the Nashville Zoo and Marley Machara for assistance with sample collection and field support. Also, thank you to Nashville Zoo veterinary staff for performing radio transmitter surgery and health checks to prepare the hellbenders for release.

## DECLARATIONS

### Authors’ contributions

CC and DMW contributed to experimental design. CC drafted the original manuscript, while DMW, JD, and LV-G reviewed and edited the original manuscript. WS, DMW, and JA provided feedback on statistical analysis, and LV-G and MG assisted with data analysis/visualization. CC, BH, WS, TM, MG, AR, SDR, PS, MF, AA, WT, EC, and DMW assisted with sample collection at the zoo or in the field. CC performed molecular wet lab work with the help of DMW and JD, and JD and MG assisted with bioinformatics. All authors reviewed and approved the final manuscript.

### Availability of data and materials

All raw sequence data are accessible in the NCBI SRA database under PRJNA1291734.

### Ethics approval

All hellbenders sampled in this study were handled with approval from Middle Tennessee State University’s Institutional Animal Care and Use Committee (IACUC 22-3002), Nashville Zoo (IACUC 23.03), Tennessee State University (IACUC WS-2021), and Tennessee Wildlife Resources Agency (TWRA; permits 5189, 1529, and 2948).

### Consent for publication

Not applicable.

