## Supplemental for "Effects of environmental setting and diet on the gut microbial ecology of eastern hellbenders (*Cryptobranchus alleganiensis alleganiensis*)"

**Supplementary Material Table of Contents**

*Supplementary Figures*

Figure S4. Taxonomic composition of the mycobiome…………………………………...4

*Supplementary Tables*

Table S1. Statistical comparison of glycerol and dry stored swabs……………………….5

Table S2. Summary of generalized linear mixed model (GLMM) results………………..6

***Supplementary Figures***
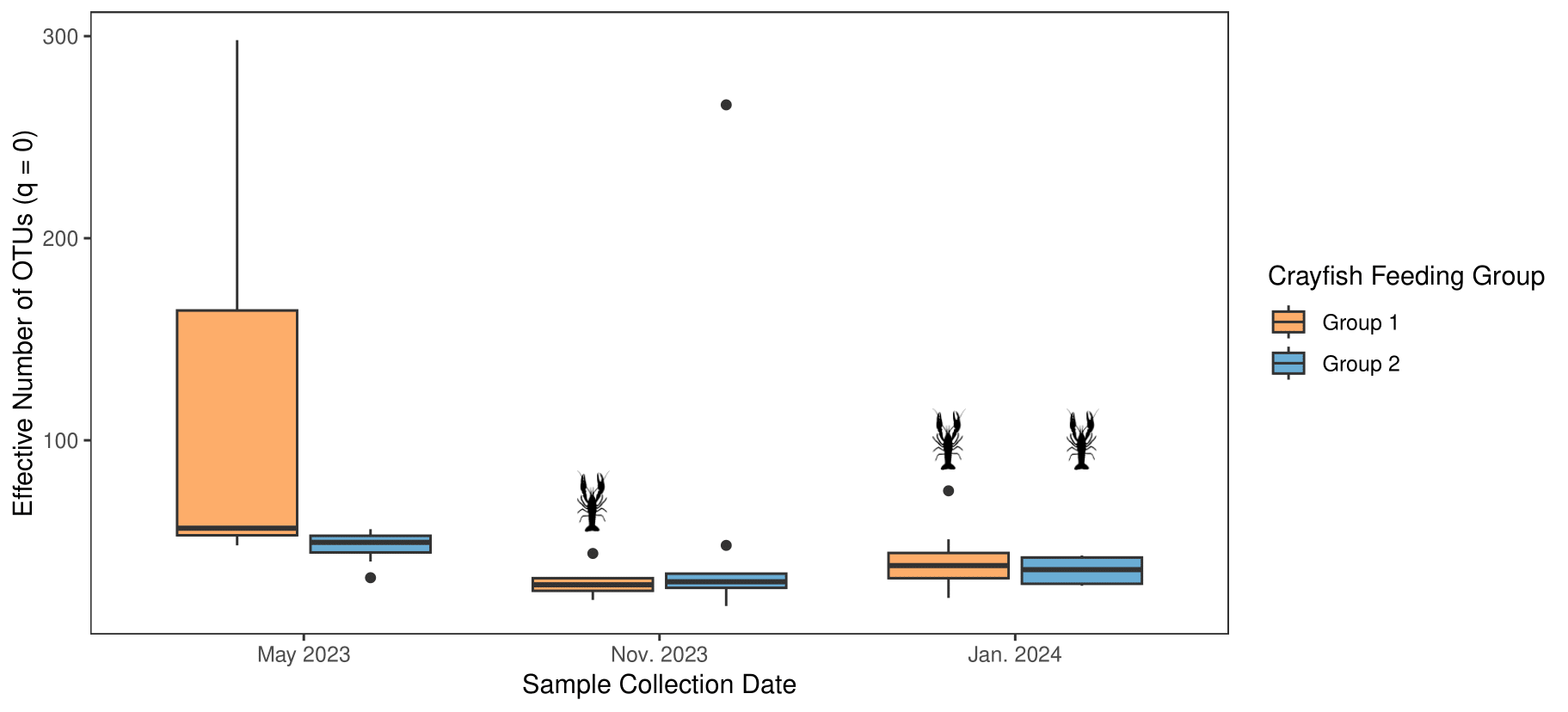

**Figure S1. Bacterial richness following crayfish feeding.** Hill-based richness (q = 0) of bacterial assemblages for two groups of male hellbenders at the Nashville Zoo that were introduced to a natural diet of crayfish across different sampling dates.

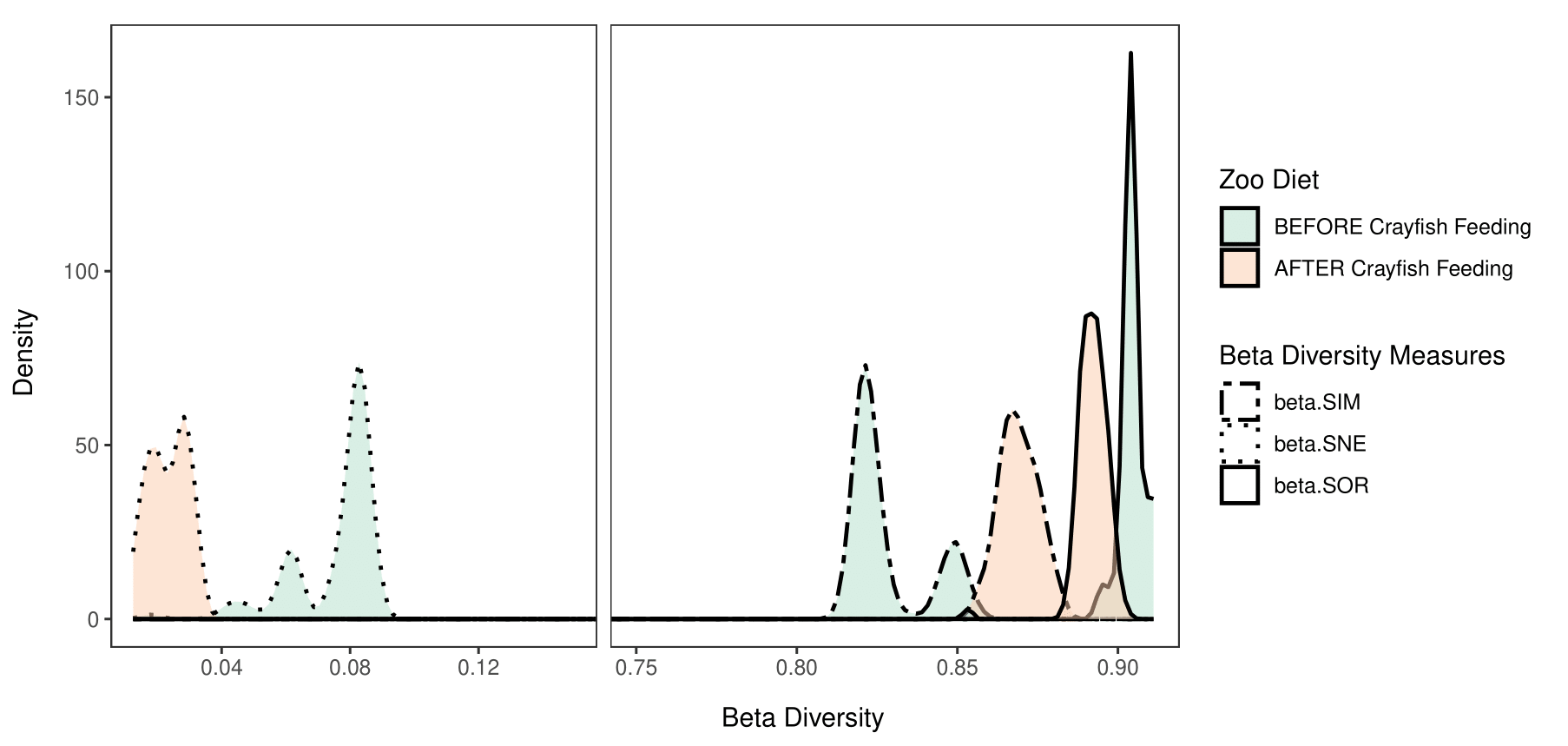

**Figure S2. Mechanisms of beta diversity following wild diet introduction.** Distribution of beta diversity components, including turnover (beta.SIM), nestedness (beta.SNE), and total beta diversity (beta.SOR), for male hellbenders exposed to a natural diet (crayfish) while at Nashville Zoo. Beta diversity values were calculated based on the Sørensen index for before and after crayfish feeding.

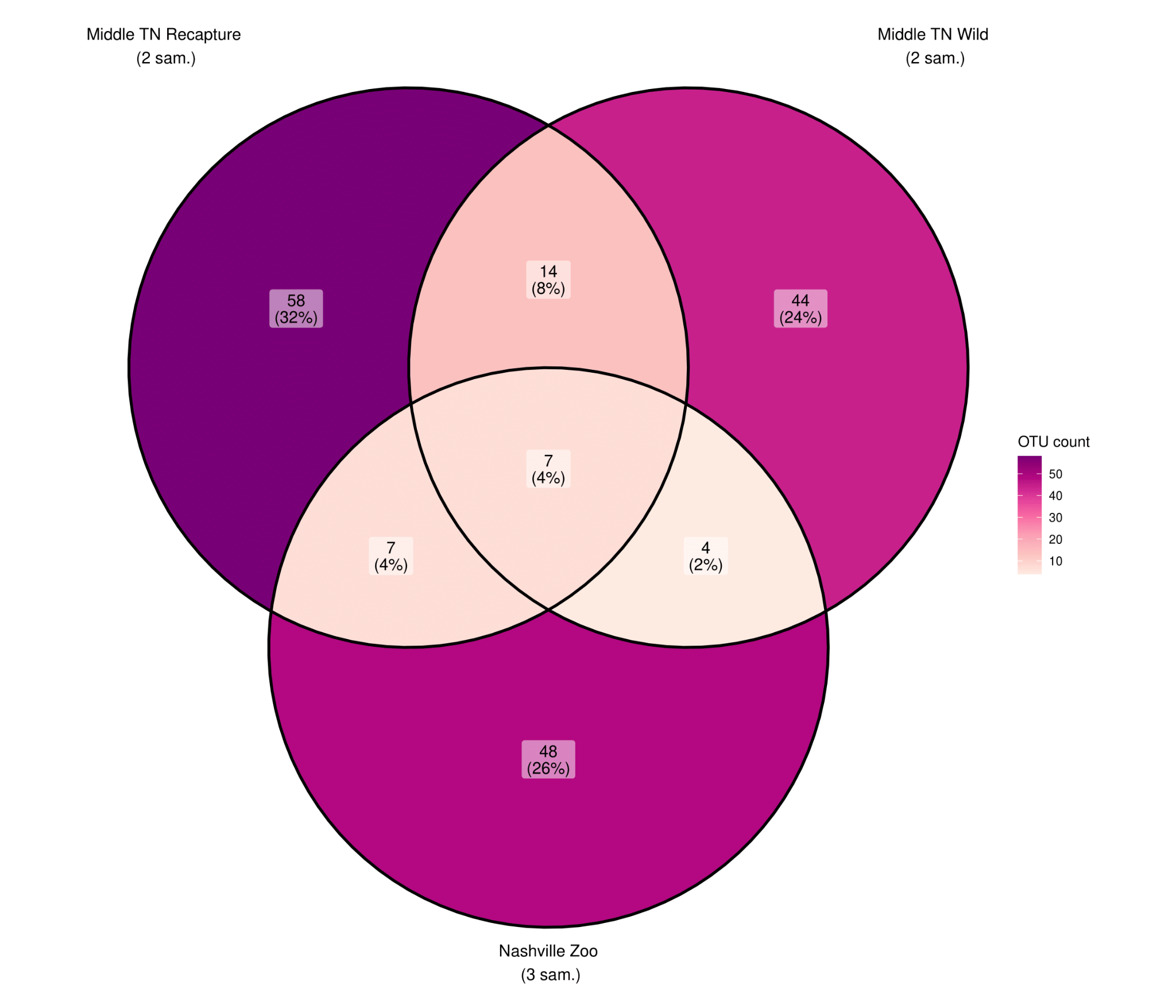

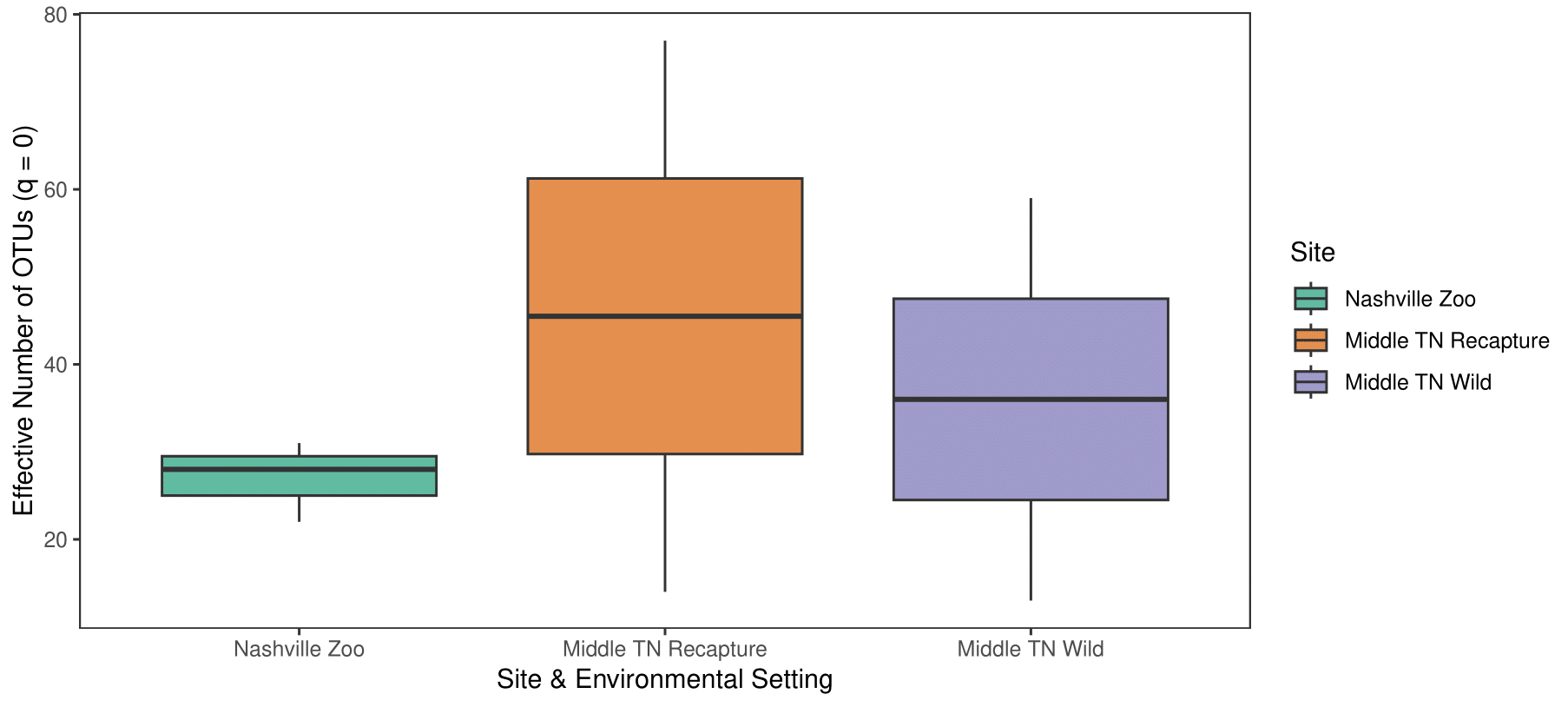

**Figure S3. Fungal richness and shared fungal diversity of the hellbender mycobiome.** a) Richness of gut fungal communities for zoo and wild-caught hellbenders, as calculated via Hill numbers (q = 0). b) Venn diagram representing the total number of shared and unique OTUs in the fungal assemblages of zoo and wild hellbenders.

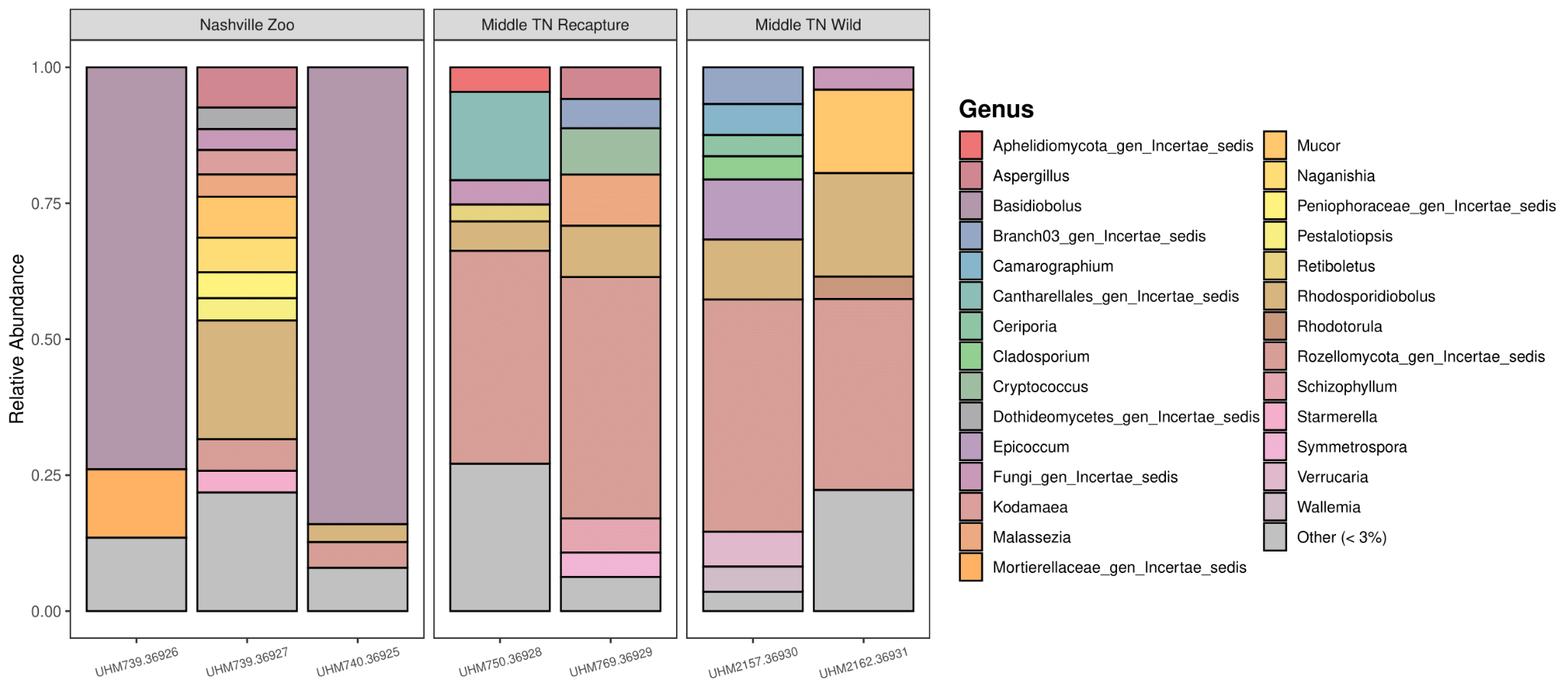

**Figure S4. Taxonomic composition of the mycobiome.** Relative abundance of fungal genera present in fecal samples collected from zoo-raised and wild-caught hellbenders. Bars represent individual hellbenders. Low abundance fungal taxa that represented less than 3% relative abundance were categorized as Other.

***Supplementary Tables***

**Table S1.** **Statistical comparison of glycerol and dry stored swabs.** Chosen statistical tests represent measures of both alpha and beta diversity.

|  | t | df | p-value | n |
| --- | --- | --- | --- | --- |
| Paired t-test (comparing bacterial richness) | -0.35319 | 5 | 0.7383 | n = 6 dry swabs  n = 6 glycerol swabs |
|  | F_1,15_ | R^2^ | Pr(>F) | n |
| Bray-Curtis PERMANOVA  (with sample storage medium as the fixed factor) | 1.9592 | 0.11552 | 0.064 | n = 11 dry swabs  n = 6 glycerol swabs |
|  | F_1,15_ | R^2^ | Pr(>F) | n |
| Raup-Crick PERMANOVA (with sample storage medium as the fixed factor) | 9291.3 | 0.99839 | 0.106 | n = 11 dry swabs  n = 6 glycerol swabs |

**Table S2. Summary of generalized linear mixed model (GLMM) results.** Results represent models of best fit as determined by comparison of AICc values.

| Response variable | Fixed effects | Estimate | Standard error | *z* value | *P* (> \|*z*\|) | Marginal R^2^ |
| --- | --- | --- | --- | --- | --- | --- |
| Bacterial richness (q = 0) for zoo and wild-caught hellbenders | (Intercept) | 4.266 | 0.439 | 9.725 | < 2e-16*** | 0.092 |
|  | Environmental setting (wild as reference category) | -0.744 | 0.286 | -2.597 | 0.009** |  |
|  | Total length (TL) | 0.008 | 0.012 | 0.691 | 0.490 |  |
| Bacterial richness for Nashville Zoo male hellbenders during wild diet manipulation | (Intercept) | 4.526 | 0.398 | 11.374 | < 2e-16*** | 0.447 |
|  | Sample collection date | -0.696 | 0.107 | -6.508 | 7.59e-11*** |  |
|  | Crayfish feeding group (Group 1 as reference category) | -0.772 | 0.253 | -3.047 | 0.002** |  |
|  | Scaled mass index (SMI) | 0.004 | 0.002 | 2.305 | 0.021* |  |
|  | Sample collection date * crayfish feeding group | 0.306 | 0.085 | 3.599 | 0.0003*** |  |
| Bacterial richness for all sampled Nashville Zoo hellbenders | (Intercept) | 2.799 | 0.845 | 3.312 | 0.0009*** | 0.030 |
|  | Age at sample collection | 0.005 | 0.139 | 0.034 | 0.973 |  |
|  | Sex (females as reference category) | -0.125 | 0.196 | -0.638 | 0.523 |  |
|  | Scaled mass index (SMI) | 0.005 | 0.002 | 2.263 | 0.024* |  |
